# The molecular link between auxin and ROS-Mediated polar root hair growth

**DOI:** 10.1101/116517

**Authors:** Silvina Mangano, Silvina Paola Denita-Juarez, Hee-Seung Choi, Eliana Marzol, Youra Hwang, Philippe Ranocha, Silvia Melina Velasquez, Cecilia Borassi, María Laura Barberini, Ariel Alejandro Aptekmann, Jorge Prometeo Muschietti, Alejandro Daniel Nadra, Christophe Dunand, Hyung-Taeg Cho, José Manuel Estevez

**Affiliations:** Fundación Instituto Leloir and IIBBA-CONICET. Av. Patricia Argentinas 435, Buenos Aires C1405BWE, Argentina; Department of Biological Sciences, Seoul National University, Seoul 151-742, Korea; Université de Toulouse, UPS, UMR 5546, Laboratoire de Recherche en Sciences Végétales, BP 42617, F-31326 CNRS, UMR 5546 Castanet-Tolosan, France; Instituto de Investigaciones en Ingeniería Gené tica y Biología Molecular, Dr. Héctor Torres (INGEBI-CONICET), Vuelta de Obligado 2490, Buenos Aires 1428, Argentina; Departamento de Qímica Biológica, IQUIBICEN-CONICET, Facultad de Ciencias Exactas y Naturales, Universidad de Buenos Aires, Intendente Guiraldes 2160, Ciudad Universitaria, Pabelloón II, Buenos Aires C1428EGA, Argentina; Departamento de Biodiversidad y Biologiía Experimental, Facultad de Ciencias Exactas y Naturales, Universidad de Buenos Aires, Intendente Güiraldes 2160, Ciudad Universitaria, Pabelloón II, Buenos Aires C1428EGA, Argentina; Instituto de Fisiologiía, Biologiía Molecular y Neurociencias, IFIByNE-CONICET, Facultad de Ciencias Exactas y Naturales, Universidad de Buenos Aires, Intendente Güiraldes 2160, Ciudad Universitaria, Pabelloón II, Buenos Aires C1428EGA, Argentina

## Abstract

Root hair polar growth is endogenously controlled by auxin and sustained by oscillating levels of reactive oxygen species (ROS). These cells extend several hundred-fold their original size toward signals important for plant survival. Although their final cell size is of fundamental importance, the molecular mechanisms that control it remain largely unknown. Here, we show that ROS production is controlled by the transcription factors RSL4, which in turn is transcriptionally regulated by auxin through several Auxin Responsive Factors (ARFs). In this manner, auxin controls ROS-mediated polar growth by activating RSL4, which then upregulates the expression of genes encoding NADPH oxidases (also known as RBOHs, RESPIRATORY BURST OXIDASE HOMOLOG proteins) and Class-III Peroxidases (PER), which catalyse ROS production. Chemical or genetic interference with the ROS balance or peroxidase activity affect root hair final cell size. Overall, our findings establish a molecular link between auxin regulated ARFs-RSL4 and ROS-mediated polar root hair growth.

**Significance Statement:** Tip-growing root hairs are excellent model systems to decipher the molecular mechanism underlying reactive oxygen species (ROS)-mediated cell elongation. Root hairs are able to expand in response to external signals, increasing several hundred-fold their original size, which is important for survival of the plant. Although their final cell size is of fundamental importance, the molecular mechanisms that control it remain largely unknown. In this study, we propose a molecular mechanism that links the auxin-Auxin Response Factors (ARFs) module to activation of RSL4, which directly targets genes encoding ROS-producing enzymes, such as NADPH oxidases (or RBOHs) and secreted type-III peroxidases (PERs). Activation of these genes impacts apoplastic ROS homeostasis, thereby stimulating root hair cell elongation.

---

In *Arabidopsis thaliana,* root hair cells and non-hair cell layers differentiate from the epidermis in the meristematic zone of the root. Once root hair cell fate has been determined, root hairs protrude from the cell surface and represent up to 50% of the surface root area, and this process is crucial for nutrient uptake and water absorption. Root hair growth is controlled by the interplay of several proteins, including the transcription factor (TF) ROOT HAIR DEFECTIVE 6 (RHD6), a class-I RSL protein belonging to the basic helix-loop-helix (bHLH) family (1). The bHLH transcription factor RSL4 (ROOT HAIR DEFECTIVE SIX-LIKE 4), which defines the final root hair length based on its level of expression (2), is a central growth regulator acting downstream of RHD6. Other TFs also contribute to the regulation of root hair growth (3,4). Root hair cell elongation is modulated by both a wide range of environmental signals (5,6) and endogenous hormones, such as ethylene and auxin (2,5,7). However, the underlying mechanisms are unknown. The transcriptional auxin response is mediated by auxin binding to receptors of the TRANSPORT INHIBITOR RESPONSE1/AUXIN SIGNALING F-BOX (TIR1/AFB) family and its co-receptor AUXIN/INDOLE 3-ACETIC ACID (Aux/IAA). This triggers the proteasome-dependent degradation of Aux/IAA, which results in the release of the Auxin Response Factors (ARFs). The released ARFs bind to *cis*-auxin response elements (Aux-REs) in the promoters of early auxin response genes to trigger downstream responses (8). Gain-of-function mutants for several Aux/IAAs exhibit reduced root hair growth (9). On the other hand, a high degree of genetic redundancy is expected in root hairs since several ARFs are highly expressed in trichoblast cells (e.g. ARF7 and ARF19 are the two most abundant ARFs (10)). In agreement, *arf7arf19* double mutant did not show a reduction in root cell elongation (11) indicating that there are other ARFs acting downstream auxin in growing root hair cells. Auxin needs to be sensed *in situ* in hair cells to trigger cell expansion, although no molecular connection has been proposed yet between ARF and downstream TFs that control growth (Fig. S1). Downstream auxin signaling is sustained by an oscillatory feedback loop comprised of two main components, calcium ions (Ca^2+^) and reactive oxygen species (ROS). High levels of cytoplasmic Ca^2+^ (_cyt_Ca^2+^) trigger ROS production by RBOHs (RESPIRATORY BURST OXIDASE HOMOLOG proteins) and high levels of _apo_ROS activate unknown Ca^2+^-permeable channels that promote Ca^2+^ influx into the cytoplasm (12,13). Plant RBOHs are plasma membrane-localized NADPH oxidases that produce apoplastic superoxide ion (O_2_^−^), which is mostly converted chemically or enzymatically into hydrogen peroxide (H_2_O_2_). *Arabidopsis* RBOHs (AtRBOHA-J) are associated with diverse ROS-related growth responses (14,15,16); for instance, RBOHC promotes root hair budding (17,18) and RBOHH,J is required for proper pollen tube growth (19). In addition, the pool of _apo_ROS can be modulated by the activity of Class-III peroxidases (PER) (20). The mechanism that links auxin to ROS-mediated cell growth is unclear (Fig. S1). Here, we propose a molecular mechanism in which endogenous auxin activates several ARFs to upregulate the expression of *RSL4,* which controls ROS-mediated polar root hair growth by promoting the expression and activity of RBOH and PER proteins. On the other hand, we were not able to detect an auxin-mediated non-transcriptional fast response on ROS production in root hair cells.

## RESULTS AND DISCUSSION

### Several ARFs-mediate RSL4 activation linking auxin-dependent polar growth to ROS production in root hair cells

To establish if ROS-production is linked to auxin-dependent polar growth, we analyzed root hair growth and ROS levels in the presence of exogenously supplied auxin (5 μM IAA, for indole 3-acetic acid). Auxin enhances root hair growth (Fig. 1A; SI Appendix, Fig. S2A) and triggers a slightly higher ROS levels in root hairs longer than 300 μ m (SI Appendix, Fig. S2B). Next, we monitored cytoplasmic H_2_O_2_ (_cyt_H_2_O_2_) levels using HyPer, which reacts with H_2_O_2_ when an external _apo_H_2_O_2_ was applied in the growing tip (SI Appendix, Fig. S3A). H_2_O_2_ levels in auxin treated roots maintain a moderate HyPer signal in root hairs longer than 700 μm while there is no root hair development in the non-treated ones that reaches these cell lengths (SI Appendix, Fig. S3B). Hence, the maintenance of _cyt_ROS levels is a reliable response to auxin in tip-growing root hairs. Auxin triggers polar root hair growth via an unknown mechanism involving RSL4 activation (5). First, we analyzed whether both RSL2 and RSL4 directly regulates ROS-production to trigger root hair growth. The *rsl2-1 rsl4-1* double mutant showed highly reduced ROS levels, while slightly higher ROS levels were detected in the RSL4-overexpressing line (RSL4^OE^) than in the wild type (Wt) during late stages of root hair development (≥ 400 μ m in length) (Fig. 1B,C). Therefore, both active RSL2 and RSL4 stimulate ROS production (Fig. 1C and SI Appendix, Fig. S2D). We then tested if auxin triggers ROS production in an RSL2-RSL4-dependent manner. Auxin partially stimulate root hair growth and ROS-production in each of *rsl4-1* and *rsl2-1* null mutants but failed in *rsl2-1 rsl4-1* double mutant (Fig. 1A,B and SI Appendix, Fig. S1C,D). Mutants for other reported TFs involved in root hair growth, such as *mamyb*(3) and *hdg11*(4), were treated with auxin, but normal responses were observed, suggesting that these TFs are not involved in auxin activation (SI Appendix, Fig. S4). In addition, the RSL4^OE^ line was less sensitive to the RBOH inhibitor VAS2870 (VAS) with respect to cell growth-linked ROS production (SI Appendix, Fig. S5A,B). Overall, these findings highlight the key role of both RSL2-RSL4 although RSL4 with a more protagonist role as main transcriptional activators of the ROS-related auxin-response in root hairs. Then, we decided to focus on how auxin activates *RSL4* expression.

**Fig. 1.**
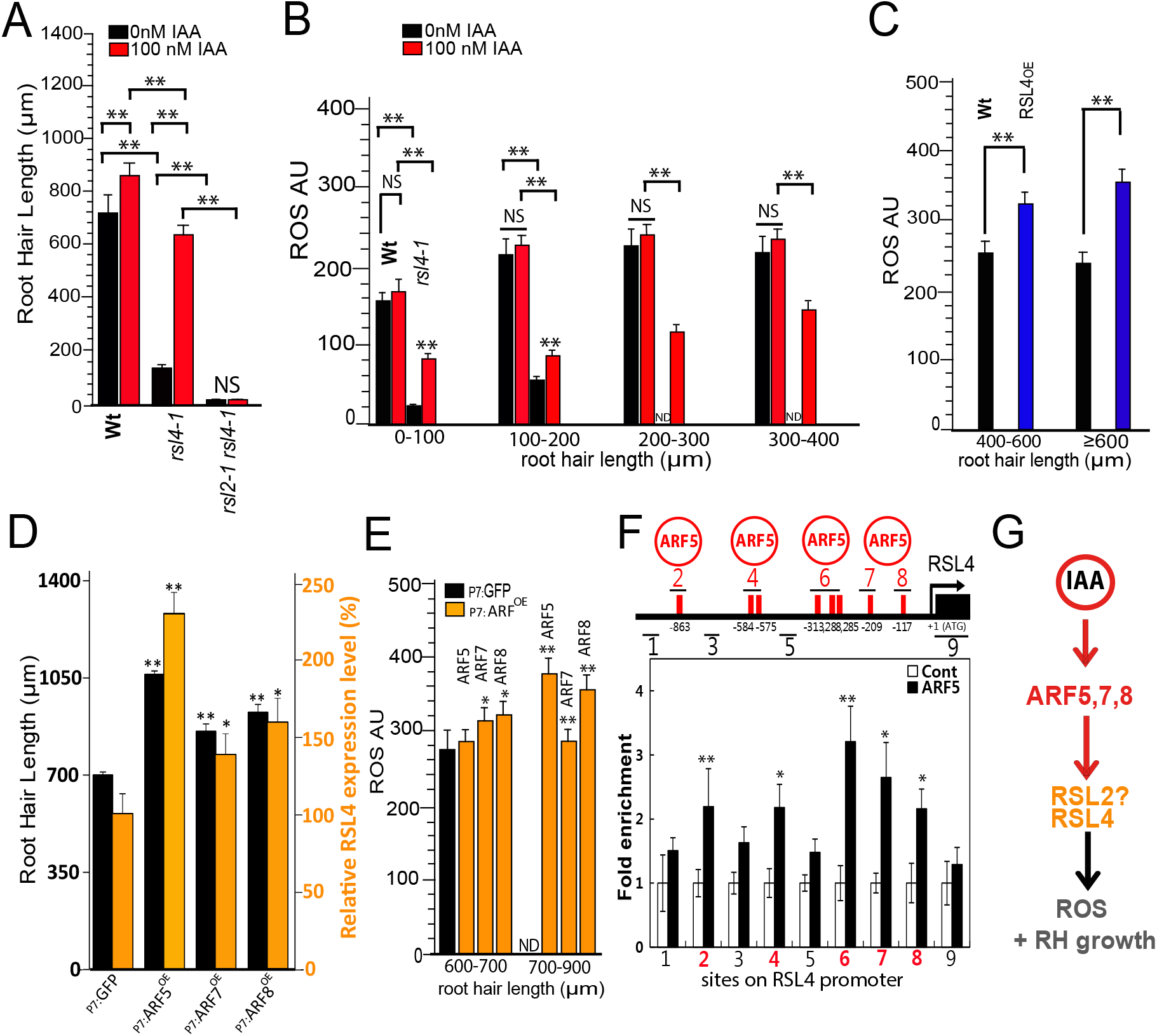
ARFs-mediated RSL4 activation links auxin-dependent polar root hair growth to ROS production. **A**, Root hair length (mean ± SD) in Wt Col-0, *rsl4-1* and *rsl2-1 rsl4-2* treated with 100nM IAA. **B,** Effect of auxin (100nM IAA, indole 3-acetic acid) mediated by RSL4 on ROS production. ROS signal analysis with IAA treatment of Wt Col-0 and *rsl4-1* mutant in the early stages of root hair development. **C**, ROS in Wt Col-0 and RSL4^OE^ (RSL4 overexpressor) in later stages of root hair growth. **D**, Effect of EXP7 (EXPANSIN 7)-driven ARF5 (_P7:_ARF5^OE^), ARF7 (_P7_:ARF7^OE^), and ARF8 (_P7_:ARF8^OE^) overexpression on polar root hair growth (black bars) and RSL4 expression (orange bars; mean ± SD). **E**, ROS in EXP7-GFP/Wt Col-0 (_P7:_GFP as control), and EXP7 driven ARF5,7,8 (_P7:_ARF5,7,8^OE^) overexpressor in the later stages of root hair development. **F**, Chromatin immuno-precipitation (ChIP) analysis showing ARF5-binding to auxin response elements (AuxREs) on the RSL4 promoter region. The RSL4 promoter region (pRSL4); the relative positions of AuxREs (red bars) and ChIP-PCR regions (lines numbered #1-9) are indicated. On the right, enrichment fold of ARF5:GFP in ChIP-PCR on each region shown. Cont (Control, pMDC7-empty line); ARF5 (pMDC7:ARF5-GFP; dexamethasone-inducible ARF5-GFP line). The values are relative to each Cont value and significantly different (^**^, P<0.001; ^*^, P<0.005; t-test) from each Cont value. **G**, Proposed events from IAA signal mediated by ARF5 activation of *RSL4* expression to ROS-linked root hair (RH) growth. In Figs. 1a-e, *P*-value of one-way ANOVA, (**) P<0.001, (*) P<0.01. NS= not significant different. ND= Not detected. In A-E Error bars indicate ±SD from biological replicates.

Next, we tested whether ARFs regulate root hair growth by upregulating RSL4 expression and ROS-production. Several ARFs like ARF5, ARF7, ARF8, and ARF19 are expressed in the meristematic and elongation zone of the root as well as in trichoblast cells and developing root hairs (Fig. S6A-B). Based on the lack of phenotype for *arf7arf19* mutant (11), all suggest a high degree of functional redundancy between these ARFs possibly involved in root hair growth. To overcome this, we then generated lines overexpressing these ARFs under the control of the strong root hair specific promoter EXPA7 (_P7_:ARF; SI Appendix, Fig. S6B). Root hair growth was enhanced in three overexpression lines tested, especially in ARF5 line (Fig. 1D; SI Appendix, Fig. S6C). Furthermore, RSL4 expression was elevated (Fig. 1D) and ROS levels were maintained high in the root hair cells of these lines during late developmental stages (Fig. 1E). Given that the region upstream of RSL4 contains eight ARF-responsive elements (Aux-RE) (Fig. 1F; SI Appendix, Table S2), we speculate that these ARFs would be able to bind to the RSL4 promoter and trigger its expression. To test this possibility, we carried out chromatin immunoprecipitation (ChIP) analyses (Fig. 1F) using an estradiol-inducible version of ARF5-GFP since it was the most efficient ARF tested to upregules *RSL4* expression. ChIP of ARF5 showed that the fold enrichment levels are significantly higher in the AuxRE regions than in the control (Fig. 1F). These data indicate that ARF5 binds to Aux-REs in the RSL4 promoter region and positively regulates its expression. It is highly possible that ARF7 and ARF8 also bind and regulates *RSL4* expression. Previously, it was shown that Aux/IAA14 *slr1-1* resistant mutant lacked root hairs (9) suggesting a strong inhibition on these ARFs. To test if a single ARF is able to overcome this inhibition, we selected ARF 19 since it is one of the ARFs most highly expressed in root hair cells (10). High levels of ARF19 (ARF19 ^OE^, overexpression) were transgenically expressed in the Aux/IAA14 *slr1-1* root hair-less background. ARF19^OE^ was able to completely rescue the *slr1-1* root hair phenotype and normalized ROS level (SI Appendix, Fig. S7A,B). As expected, ARF19 nuclear expression is highly responsive to auxin and it is localized in root epidermal cells close to the root hair development zone (SI Appendix, Fig. S7C,D). Together, these results and previous reports (9-11) suggest that high auxin levels in trichoblast cells would release several ARFs (e.g. ARF5, ARF7, ARF8, and ARF19) from its repressors Aux/IAAs, to directly control RSL4 expression (and possibly RSL2), triggering ROS-mediated root hair elongation (Fig. 1F). Further experiments are required to establish how these ARFs are regulated to act in an articulate manner to trigger the transcriptional response during cell growth.

## RSL4-regulated *RBOHC, J* as well as *RBOHH* drive ROS-mediated root hair growth

Besides RBOHC, two other RBOH groups exist; one group clusters RBOHC and is comprised of RBOHE,F (SI Appendix, Fig. S8A) and the other was previously implicated in polar-growth of pollen tubes^18^ and is composed by RBOHH,J. To identify which of these RBOHs also contribute to ROS-linked to tip-growth, we isolated *rboh* mutants (SI Appendix, Table S1; Fig. S8B) and screened them for abnormal cell expansion linked to deficient ROS production (Fig. 2A,B). Only *rbohc-1* and *c-2* showed short root hairs, whereas *rbohh-1* and *h-3* developed slightly less elongated cells (SI Appendix, Fig. S8C,D). When double *rboh* mutants were analysed, only *rbohh,j* and mutants containing *rbohc* showed a clear root hair growth reduction of up to 40% (Fig. 2A). In agreement with this, *rbohc* mutants showed highly reduced ROS levels, while double *rbohhj* mutants presented moderate to low ROS levels (Fig. 2B). We then used the HyPer H_2_O_2_ sensor to test if the RBOH inhibitor VAS could limit the level of _cyt_H_2_O_2_ in the growing root hair tip. The level of _cyt_H_2_O_2_ was significantly diminished after VAS treatment (Fig. 2C) and _cyt_H_2_O_2_ signatures were drastically modified in *rbohc-1* root hairs (Fig. 2D). Exogenous H_2_O_2_ treatment failed to rescue *noxc-1* root hair growth defect (Fig S9A) implying that a complex and fine-tuned ROS regulation would be required to trigger growth. Together, these data suggest that although RBOHC is the main RBOH involved in ROS-mediated root hair growth, as reported previously (12,16), RBOHHJ are also important ROS-producing enzymes in this process.

**Fig. 2.**
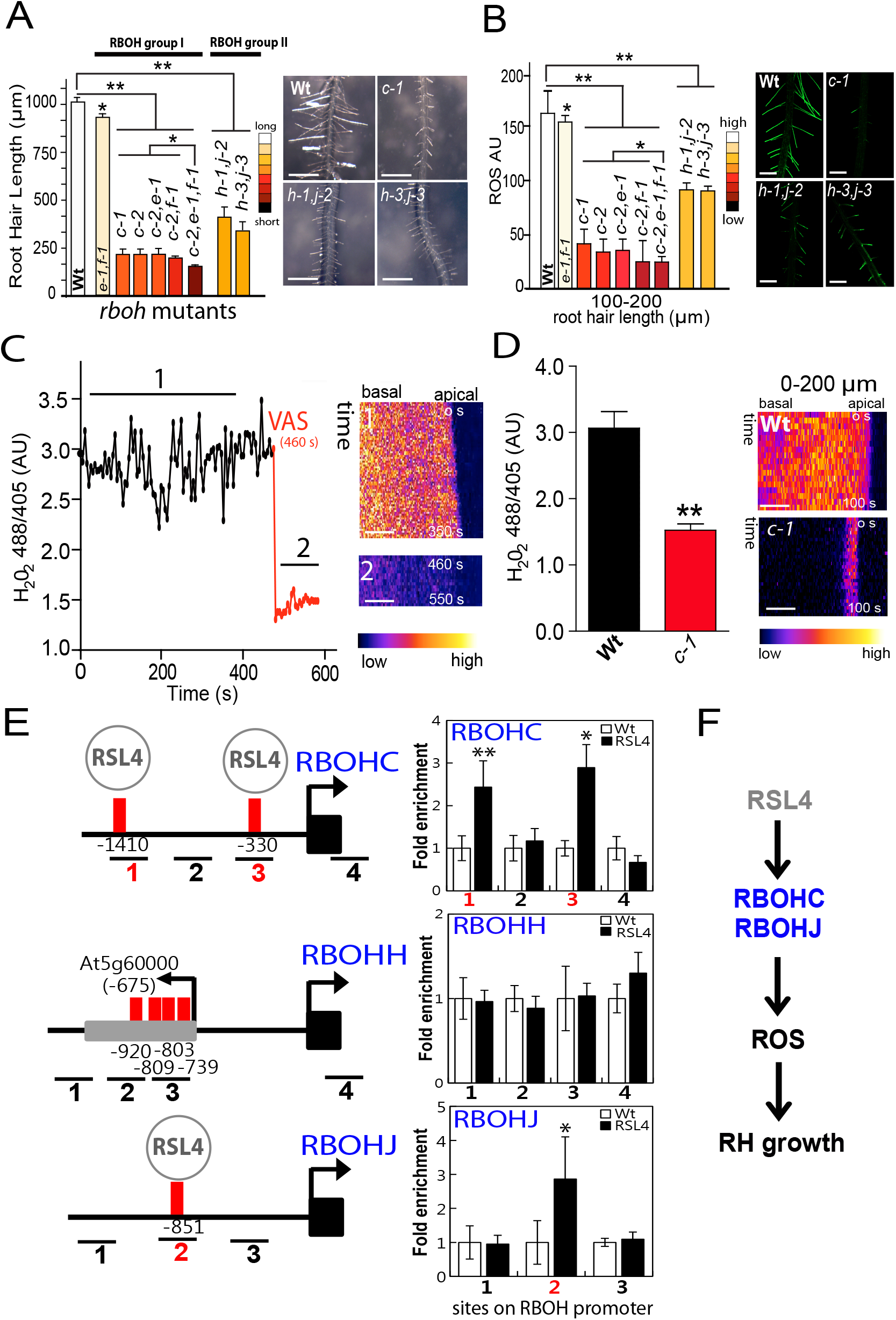
RSL4-regulated *RBOHC, J* as well as *RBOHH* drive ROS-mediated root hair growth. **A**, Root hair phenotype in Wt Col-0 and *rboh* mutants (right panel). Scale bar = 600 μm. Root hair length (mean ± SD, n roots= 30) in *rboh* mutants (left panel). **B**, Total ROS generated by oxidation of H_2_DCF-DA in early stages of root hair development in *rboh* mutants. AU= fluorescent arbitrary units. On the right, total ROS H_2_DCF-DA staining in roots of Wt Col- 0 and *rboh* mutants. Scale bar = 600 pm. **C**, _cyt_H_2_O_2_ levels in VAS-treated Wt Col-0 root hairs expressing HyPer sensor. _cyt_H_2_O_2_ levels are based on the ratio 488/405 nm of HyPer biosensor at the root hair tip over 600 s. On the right, selected kymographs resulting of this analysis only for root hairs of □200 μm in length. Scale bar = 5 μ m. **D**, _cyt_H_2_O_2_ average levels in *rbohc-1* and Wt Col-0 root hairs expressing HyPer. Average _cy_tH_2_O_2_ levels are based on the 488/405 nm ratio of HyPer at the root hair tip in root hairs of 0-200 pm in length. On the right, selected kymographs. Scale bar = 5 μ m. **E**, Promoter regions of ROS-related genes *RBOHC, H,* and *J* as targets of the RSL4 response element (RHE; red bar) and ChIP-PCR regions (lines numbered #1-4). Neighboring gene exons are indicated as gray boxes. Enrichment fold of RSL4-GFP in ChIP-PCR on each region shown. Wild type (Wt) and _pRSL4_:RSL4-GFP (RSL4). Error bars indicate ±SD for biological replicates (n=2). The values are relative to each Cont value. Significantly different (**, P<0.01; *, P<0.05; t-test) from the Cont value. F, Proposed events from RSL4 activation of RBOHC,J to ROS-mediated polar root hair (RH) growth. In Figs. 2a,b,d *P*-value of one-way ANOVA, (**) P<0.001, (*) P<0.01.

Based on an analysis of RSL4-regulated genes (5), a putative RSL4 responsive element (RSL4-RE) with the sequence TN_6_CA[CT]G[TA] was identified that is highly similar to the Root Hair *cis*-Element (RHE) TN_5-6_CACG[TA]^20^. The promoter regions of *RBOHC, H,* and *J* genes include several RHEs suggesting that these three RBOHs could be direct targets of the RSL4 TF (SI Appendix, Table S2). Using ChIP analysis, we examined whether RSL4-GFP could bind to the promoter regions of RBOHC,H,J *in vivo.* Except for RBOHH gene, RSL4 bound to at least one RHE region in RBOHC and RBOHJ promoters (Fig. 2E). This result suggests that RSL4 directly regulates these two ROS-related genes to generate the ROS required for tip growth.

Why are three RBOH proteins needed to trigger polar-growth in a single cell?. The simplest hypothesis is that these RBOHs act sequentially. Low levels of ROS were detected when the root hair first emerged from the root, but the ROS signal was up to two-fold greater in slightly longer Wt Col-0 root hairs (of 200-250 μm in length) (SI Appendix, Fig. S9,E). At a comparable stage, roots hairs of the *rbohc-1* mutant had very low levels of _cyt_ROS and those of the *rbohh,j* double mutant had only slightly lower ROS levels than did those of Wt Col-0. In later growth stages, _cyt_ROS levels were drastically reduced in the double *rbohhj* mutant. This suggests that RBOHC is the first RBOH to produce ROS, and that RBOHC functionally overlaps with RBOHH,J. Finally, RBOHHJ produced ROS that was linked to root hair tip elongation at later developmental stages (□200 pm in length; fig. S9E). Next, we investigated whether ROS is required for root hair initiation by subjecting Wt Col-0 roots to VAS. The concentration of VAS needed to inhibit root hair growth by half (IC_50_) was ~7.5 μM (SI Appendix, Fig. S9B), which is similar to ROS VAS IC_50_ (~7.0 μM) (SI Appendix, Fig. S9C). The *rbohc-1* mutant (12) showed very short root hairs with up to 150 μm in length (Fig. S9B) but still produces ~30% of ROS compared to Wt root hairs (Fig. 2D; SI Appendix, Fig. S9C) indicating the presence of a residual ROS production (17). Since ROS homeostasis could be modified by the activity of PER, we treated Wt and *rbohc-1* mutant roots with the PER inhibitor SHAM (salicylhydroxamic acid) (22), at concentrations of up to 100 μ M. SHAM treatment resulted in a significant reduction in total ROS in *rbohc-1* in comparison with non-treated *rbohc-1,* although root hair initiation was not abolished (SI Appendix, Fig. S10A,B). In an analysis using HyPer, 100 μM SHAM completely abolished the H_2_O_2_ signal in the whole root (SI Appendix, Fig. S11A,B) confirming that PERs also contribute to ROS production. This implies that, at least in *Arabidopsis thaliana,* initial development of the root hair tip is not affected by highly reduced ROS levels.

## RSL4-regulated *PERs* affect ROS homeostasis and polar root hair growth

Next, co-expression analysis revealed that *RSL4* is highly co-regulated at the transcriptional level with several *PER* genes, most of which exhibited root hair expression (SI Appendix, Fig. S12A,B) (5). We then analyzed the corresponding T-DNA *per* mutants (SI Appendix, Table S1; Fig. S12C). The single mutant *per1-2* and double mutant *per44-2,73-3* (all null mutants except for *per1-2* knock-down; SI Appendix, Fig. S12D) showed mild phenotypes (short root hairs), with up to ~30% reductions in ROS levels (Fig. 3A,B). This suggests that there is genetic redundancy between these PERs in the regulation of ROS homeostasis. The levels of _cyt_H_2_O_2_, as visualized with HyPer, were reduced in root hairs treated with 75 μ M SHAM (Fig. 3C), and this result was confirmed using 2′,7′-dichlorodihydrofluorescein diacetate (H_2_DCF-DA) probe (SI Appendix, Fig. S10B). In addition, peroxidase activity was significantly lower in SHAM-treated roots as well as in *per* mutants, higher in PER44 overexpressor (PER44^OE^) than in Wt non-treated ones (Fig. 3D) suggesting a direct link between peroxidase activity and ROS-homeostasis. To explore the transcriptional regulation of these *PERs* in ROS-mediated root hair growth, we analyzed the regulatory region of the genes encoding these *PERs.* We identified several putative RHE motifs (SI Appendix, Table S3), suggesting that these genes could be direct targets of RSL4, as suggested by the co-expression analysis (SI Appendix, Fig. S12A). ChIP assays revealed that RSL4 binds to the RHE of four *PERs, PER1, 44, 60,* and *73* (Fig. 3E). In agreement with this, much higher and lower levels of *PER1* and *PER73* were detected in RSL4 ^OE^/Wt and *Wt/rsl4-1* lines, respectively (5), suggesting that RSL4 promotes the expression of at least these two *PERs.* At last, higher peroxidase activity was found in _P7:_ARF5 and _P7:_ARF8 lines (where RSL4 expression is upregulated; Fig 1D) and much lower activity was detected in *rsl4-1* that lacks RSL4 (Fig. 3F) confirming that RSL4 not only directs *PER* expression at transcriptional level but also impacts on the PER activity (Fig. 3G).

**Fig. 3.**
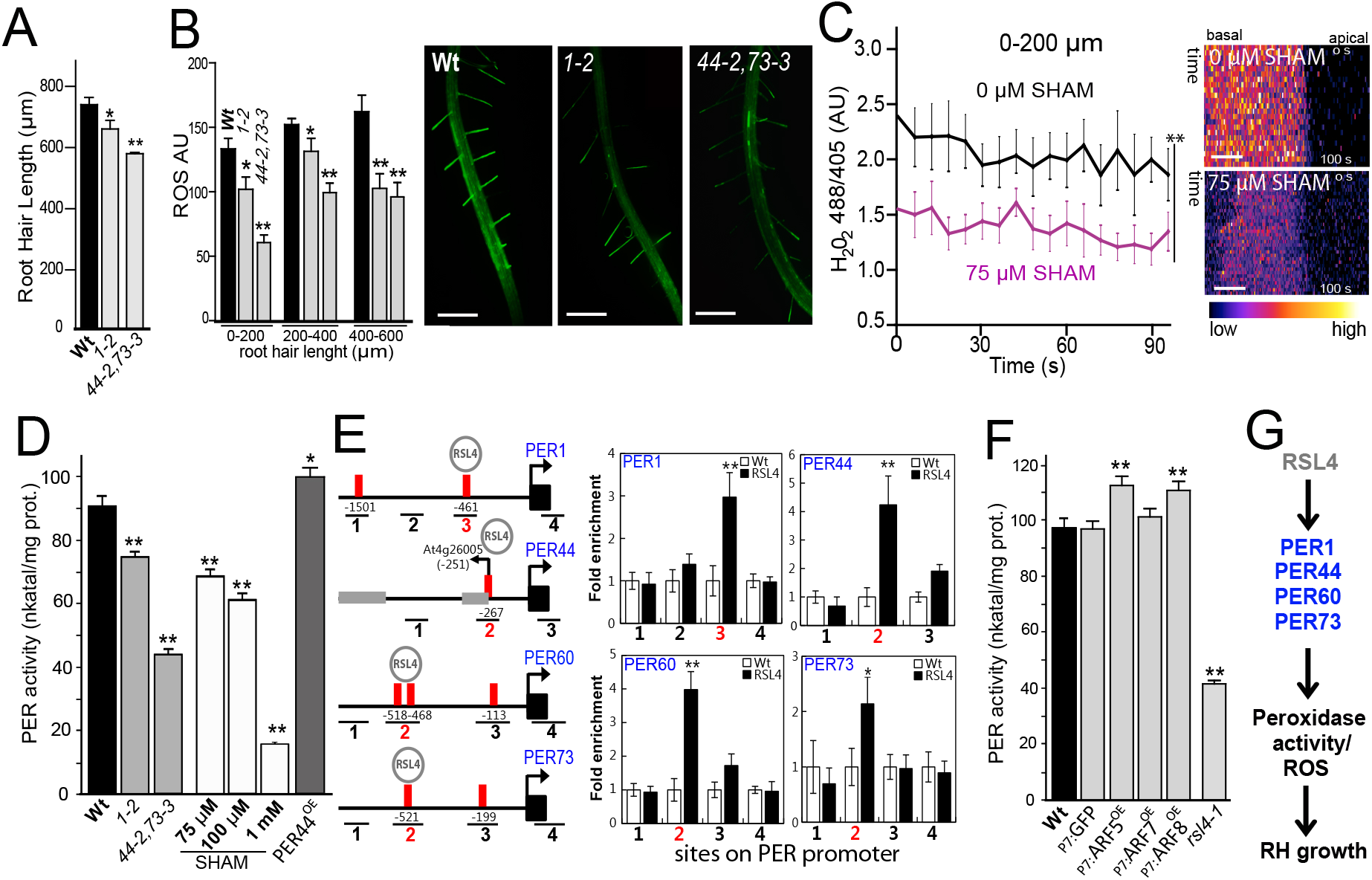
RSL4-regulated *PERs* affect ROS homeostasis and polar root hair growth. **A**, Root hair length (mean ± SD, n roots= 30) in the single mutant *per1-2* and double mutant *per44-2,73-3.* P-value of one-way ANOVA, (**) P<0.001, (*) P<0.01. **B**, ROS in root hairs of single mutant *per1-2* and double mutant *per44-2 per73-3.* On the right, ROS H_2_DCF-DA staining in roots of Wt Col-0 and *per* mutants. Scale bar = 600 μm. **C,** Average _cyt_H_2_O_2_ levels in SHAM (salicylhydroxamic acid)-treated and in non-treated Wt Col-0 root hairs expressing HyPer sensor (n=10). Average _cyt_H_2_O_2_ levels are based on the 488/405 nm ratio of HyPer at the root hair tip over 100 s. *P*-value of one-way ANOVA, (**) P<0.001. On the right, selected kymographs. Scale bar = 5 μm. D, Peroxidase activity in the roots of wild type (Wt), *per1-2, per44-2, per73-3* simple mutants, *per44-2,73-3* double mutant and on wild type root extracts with increasing concentrations of SHAM. Enzyme activity (expressed in nkatal/mg protein) was determined by a guaiacol oxidation-based assay. Values are the mean of three replicates ± SD. E, Promoter regions of ROS-related genes PER1,44,60,73 showing the RSL4 response elements (RHEs; red bar) and ChIP-PCR regions (lines numbered #1-4). Positions are relative to the start codon. Neighboring gene exons are indicated as gray boxes. Enrichment fold of RSL4:GFP in ChIP-PCR on each region shown. Wild type (Wt); RSL4 (pRSL4:GFP). Error bars indicate ±SD of biological replicates (n=2). The values are relative to each Wt value and significantly different. (**) P<0.01 (*), P<0.05; t-test. **F**, Peroxidase activity in the roots of wild type (Wt), EXP7 (EXPANSIN 7)-driven GFP (as control), ARF5 (_P7_:ARF5^OE^), ARF7 (_P7_:ARF7^OE^), and ARF8 (_p7:_ARF8^OE^) as well as in *rsl4-1* mutant. Enzyme activity (expressed in nkatal/mg protein) was determined by a guaiacol oxidation-based assay. Values are the mean of three replicates ± SD. G, Proposed events from RSL4 activation of PER1,44,60,73 to ROS-linked root hair (RH) growth. In Figs. 3a,b,d,f, P-value of one-way ANOVA, (**) P<0.001, (*) P<0.01.

We then examined if auxin treatment could compensate for the lack of RBOH and PER proteins (Fig. 4A,B). Auxin treatment partially rescued the reduced length and ROS levels of the *rbohc-1* mutant, resulting in ~30% increases in both, and almost fully rescued the defects in the *rbohh,j* and *per44-2,73-3* mutants (Fig. 4A,B). When 15 μM VAS was added along with the auxin (IAA), the *rbohc-1* mutant exhibited intermediate root hair growth and partial recovery of ROS levels (SI Appendix, Fig. S13A,B) suggesting other sources of ROS. When SHAM was added in addition to VAS (SHAM+VAS) in the presence of auxin, a low recovery of root hair growth and ROS levels was detected in *rbohc-1* and Wt root hairs (SI Appendix, Fig. S13C,D). Together, these findings confirm that auxin requires both RBOH-derived ROS and ROS produced by PER to modulate polar growth (SI Appendix, Fig. S13E). The small although detectable recovery in ROS and cell growth after auxin treatment in root hair cells (in *rbohc-1* as well as in Wt) in the presence of both inhibitors, VAS and SHAM (SI Appendix, Fig. S13C-D), would possibly suggest the existence of an unknown source of ROS coming from other apoplastic ROS-producing enzymes (e.g. oxalate oxidase, diamine oxidase, lipoxigenases, etc.) also regulated by auxin-RSL4 (and possibly RSL2) (SI Appendix, Fig. S13E) that will require further investigation.

**Fig. 4.**
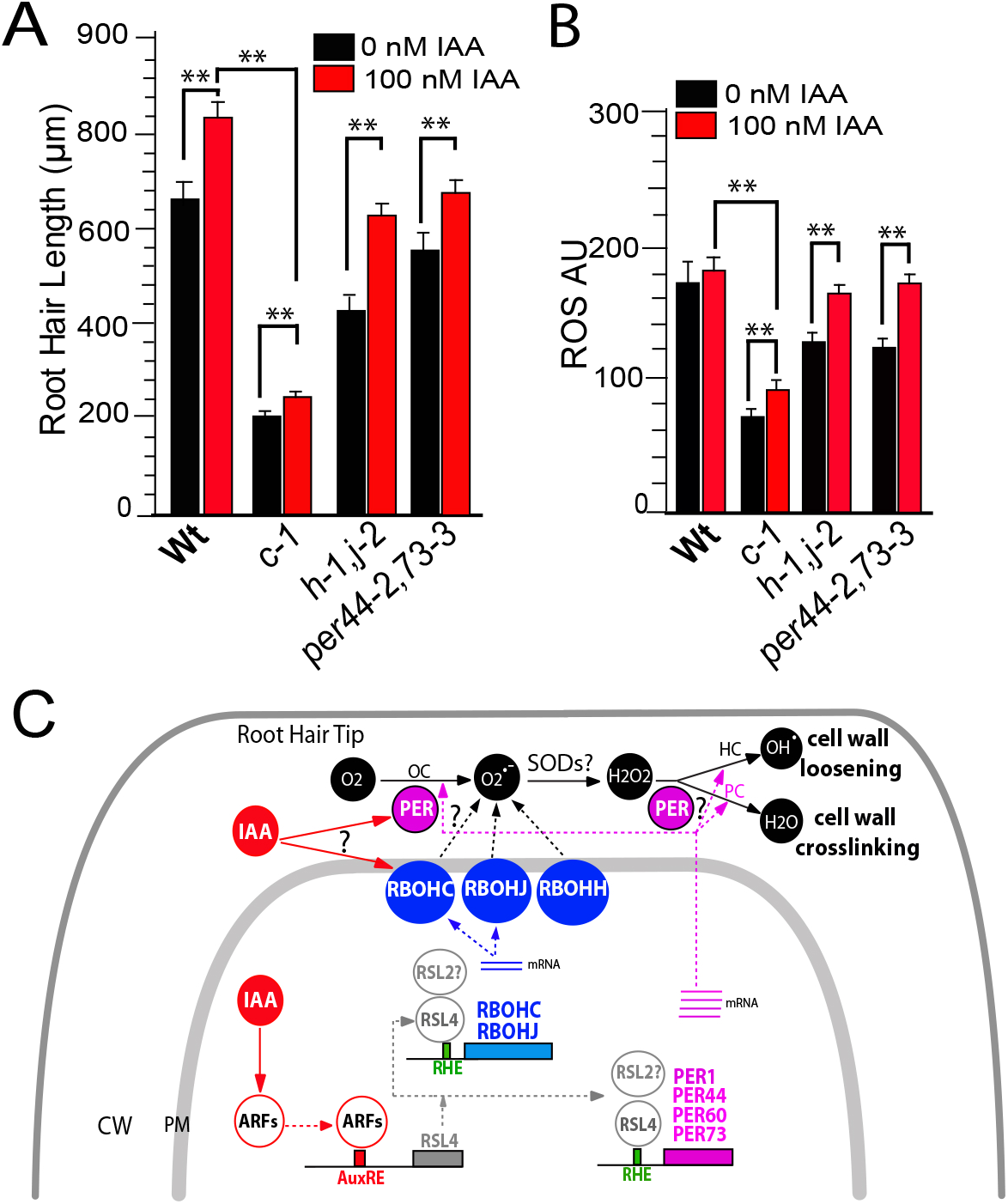
Auxin-stimulated polar growth requires apoplastic ROS. **A**, Root hair length (mean ± SD., n roots= 30) in Wt Col-0, *rboh* and *per* mutants treated or not with 100nM of IAA. **B**, ROS signal in Wt and *rboh* mutants treated or not with 100nM of IAA. AU= arbitrary units. Root hairs of 0-200 μ m in length were analyzed. C, Model of auxin-ARF5-RSL4 regulation of ROS mediated polar root hair growth. The bHLH transcription factor RSL4 is transcriptionally activated by auxin (IAA) and its expression is directly regulated by several ARFs (e.g. ARF5,7,8,19). Throughout RSL4 (and possibly RSL2), auxin activates the expression of two RBOHs (RBOHC,J) and four PERs (PER1,44,60,73) that together regulate ROS homeostasis in the apoplast (in combination with RBOHH). PERs in the presence of _apo_ROS would control cell wall loosening (in the peroxidative cycle, PC) and crosslinking reactions (in the hydroxylic cycle, HC) that impact on polar growth. PERs also contribute to the superoxide radical pool by oxidizing singlet oxygen (in the oxidative cycle, OC). Solid lines indicate activation or chemical conversion. Dash lines indicate protein translation and targeting or contribution. CW=cell wall; PM= plasma membrane. Reactive Oxygen Species (ROS): hydroxyl radical (^•^OH); superoxide ion (O_2_^•-^), hydrogen peroxide (H_2_O_2_); singlet oxygen (O_2_); superoxido dismutases (SODs). In Figs. 4a,b, *P*-value of one-way ANOVA, (**) P<0.001, (*) P<0.01. (?) means unidentified PER catalyzing the indicated reaction. Auxin response elements (AuxRE). RSL4 response element (RHE). In A-B, error bars indicate ±SD.

Finally, we tested if auxin is able to trigger a rapid and non-transcriptional response impacting on ROS production in the root hair tip (SI Appendix, Figure S14). We applied IAA in growing root hairs that express the HyPer sensor and no significant changes were detected before and after hormone treatment (SI Appendix, Fig. S14A-B). A similar result was obtained when ROS levels were measured with the dye H_2_DCF-DA (SI Appendix, Fig. S14C). Overall, our results suggest that the main auxin downstream targets are ARFs-RSL4 (and possibly RSL2) and the transcriptional activation of genes related to ROS-production and involved in polar growth (SI Appendix, Fig. S13E). We cannot exclude other direct non-transcriptionally activated targets involved in the auxin response in root hair cells but we fail to detect changes in H_2_O_2_/ROS upon auxin stimulation in our experimental conditions.

## Conclusions

A root hair is the single-cellular protrusion able to extend in response to external signals, increasing several hundred-fold its original size, which is important for survival of the plant. Final root hair size has vital physiological implications, determining the surface area/volume ratio of the whole roots exposed to the nutrient pools, thereby likely impacting nutrient uptake rates. Although the final hair size is of fundamental importance, the molecular mechanisms that control it remain largely unknown. Here, we propose a mechanism that explains how root hair tip growth works under auxin regulation of ROS homeostasis by activating several ARFs, and subsequently, upregulating the expression of the bHLH TF RSL4 (and possibly RSL2). RSL2-RSL4-dependent ROS-production is catalyzed by RBOH and PER enzymes, which facilitates polar root hair growth and defines its final cell size (Fig. 4C). Auxin is a prominent signal in trichoblast cells and is essential for proper polar root hair growth. Auxin-ARF activation of bHLH TF regulates several other plant developmental processes (25,26). Here, we have identified that auxin activates *RSL4* via several ARFs (ARF5,7,8), which binds (at least ARF5) to several Aux-REs in the RSL4 regulatory region (8). In addition, high levels of ARF19 are able to overcome a resistant Aux/IAA14 *slr1-1* to trigger ROS mediated polar growth. Recently, two dominant mutated versions of Aux/IAA7, were found to completely suppress root hair growth by down-regulating RSL4 expression (27). Based on these results, we hypothesize that IAA7-IAA14 directly represses several ARFs (ARF5,7,8,19) to control RSL4 (and RSL2) expression. Based on the mutant analysis, RSL2 would act together with RSL4 to trigger ROS responses when stimulated with auxin. Once RSL4 is active, it will trigger RSL2 expression (2,5) and both are able to control ROS homeostasis during polar root hair growth by directly regulating the expression of *RBOHs* and *PERs.* Different types of ROS affect cell growth by modulating the balance between cell proliferation and cell elongation in the root meristem (28,29), by unknown molecular mechanism. In this study, we report that, under auxin induction, RSL2-RSL4 triggers ROS-mediated cell elongation. ChIP analysis revealed that RSL4 directly regulates the expression of two *RBOHs* as well as four *PERs.* However, out of 34 putative direct targets of RSL4 reported (30), only few (e.g. PER7) were found to be associated with ROS-homeostasis. In another recent report, RSL4 was shown to bind to RHE and trigger (directly or indirectly) the expression of the smallest subset of 124 genes involved in ROS-homeostasis, cell wall synthesis and remodeling, metabolism, and signaling that are necessary and sufficient to trigger root hair growth (31). Still unknown are the RSL2 targets and how RSL2 acts in a cooperatively way with RSL4. In addition, the transcriptional mediator subunit (MED) MED15/ PHYTOCHROME AND FLOWERING TIME1 (PFT1) affects ROS levels by triggering transcriptional changes in root hair cells (32), probably via a mechanism that does not involve RSL4. Alternatively, ROS homeostasis may be influenced by other factors, independently of the RSL4 transcriptional program, that also modulate RBOH and PER activities, such as Ca^2+^ binding and posttranslational modifications (18,33). Overall, here we established a key role of RSL4 as one transcriptional activator of the ROS-related auxin-response in root hairs together with RSL2 but we cannot discard that other TFs could also act downstream auxin and independently of RSL2-RSL4 to trigger ROS-production in root hair cells (SI Appendix, Fig. S13E).

Here, we identified two RBOHs (RBOHH and J) together with the previously reported RBOHC (12) and four PERs (PER1, 44, 60, and 73) that are involved in ROS-mediated polar root hair growth in *Arabidopsis* (Fig. 4C). In agreement with our findings, a RBOH named RTH5 (for *RooT hair Less* 5) from maize with high sequence similarity to AtRBOHH,J was also linked to root hair cell expansion (34). Several of the PERs identified as contributing to ROS-mediated root hair growth, were previously found to modulate cell growth, but were not shown to be involved in ROS homeostasis (30,35). Exogenous supplied H_2_O_2_ inhibited root hair polar expansion, while treatment with ROS scavengers (e.g., ascorbic acid) caused root hair bursting (16), reinforcing the notion that _apo_ROS modulates cell growth by impacting cell wall properties. In this manner, _apo_ROS molecules (_apo_H_2_O_2_) coupled to PER activity directly affect the degree of cell wall crosslinking (14,20) by oxidizing cell wall compounds and leading to rigidification of the wall in peroxidative cycles (PC) (16). The identified PER1,44,73 directly contribute to PER activity in PC. On the other hand, _apo_ROS coupled to PER activity enhances non-enzymatic wall-loosening by producing oxygen radical species (e.g., ^•^OH) and promoting polar-growth in hydroxylic cycles (HC) (36). It is unclear how opposite effects on cell wall polymers are coordinated during polar growth (20,36). Finally, PERs also contribute to the superoxide radical (O_2_•^−^) pool by oxidizing singlet oxygen in the oxidative cycle (OC), thereby affecting _apo_H_2_O_2_ levels. Although ROS is produced in the apoplast influencing the cell wall properties during cell expansion, H_2_O_2_ is also present at high levels in the cytoplasm during polar growth (17). In agreement, H_2_O_2_ influx from the apoplast was demonstrated for several plant PIPs in heterologous systems (23,24,37) as well as in plant cells (23). Since cytosolic acidification triggers a drastic reduction in the water permeability of several PIPs, it is plausible that oscillating pH changes in growing root hairs (17) directly regulate the cycles of _cyt_H_2_O_2_ influx. Once H_2_O_2_ molecules are transported into the cytoplasm, they trigger downstream responses linked to polar growth (12,18).

## Materials and Methods

### Root hair phenotype

For quantitative analysis of root hair phenotypes in *rboh* mutants and Wt Col-0, 200 fully elongated root hairs were measured (n roots= 30) from seedlings grown on vertical plates for 10 days. Values are reported as the mean ±SD using the Image J 1.50b software. Measurements were made on images were captured with an Olympus SZX7 Zoom microscope equipped with a Q-Colors digital camera. 10 days old roots for ARF^OE^ were digitally photographed with a stereomicroscope (M205 FA, Leica, Heerbrugg, Switzerland) at a 40X magnification. The hair length of 8 to 10 consecutive hairs protruding perpendicularly from each side of the root, for a total of 16 to 20 hairs from both sides of the root.

### H_2_DCF-DA probe used to measure total ROS

Growth Arabidopsis seeds in plate with agar 1% sterile for 8 days in chamber at 25°C with a continuous light. These seedlings were incubated in darkness in a slide for 10 min with the 2′,7′-Dichlorodihydrofluorescein diacetate (H_2_DCF-DA) 50 μM at room temperature. Samples were observed with a confocal microscope equipped with 488 nm argon laser and BA510IF filter sets. It was used 10X objective; 0.30 NA; 4.7 laser intensity; 1.1 off set; 440 PMT (for highest ROS levels) 480 PMT (for ROS media) and 3 gain. Images were taken scanning XZY with a 2 um between focal planes. Images were analyzed using ImageJ. To measure ROS highest levels, a circular ROI (r=2.5) was taken in the zone of the root hair with highest intensities. To measure ROS mean the total area of the root hair was taken. Pharmacological treatments were carried out with a combination of the following reagents: 100 μM and 5 μM IAA, 15 μM of VAS2870, 100 μM SHAM. In the case of the short time experiment with IAA, 8 days old Wt plants were incubated with 100 nM IAA for 3 minutes and co-incubated 7 minutes with 100 nM IAA and 50 μ M H_2_DCFDA at room temperature. We washed the sample with a MS0.5X solution and the image acquisition was made with 10X objective and 400ms of exposure-time in an **epifluorescence** microscope (Zeiss, Imager A2). As control we incubated 10-days old Wt plants with MS 0.5X solution for 3 minutes and coincubated 7 minutes with MS0.5X and H_2_DCFDA 50 μ M at room temperature. To measure ROS levels a circular ROI (r=2.5) was taken in the tip of the root hair. Values are reported as the mean ±SD using the Image J 1.50b software.

## Acknowledgements

We would like to thank T. Beeckman, A. Costa, L. Cárdenas, I. De Smet, L. Dolan, N. Geldner, U. Grossniklaus, T. Hamann, J.-Y. Kim, J. Kleine-Vehn, and D. Weijers for materials. We thank A. R. Kornbliht for advice and K. Farquharson (Plant Editors) for English copy edition. J.M.E and H.-T.C. laboratories for discussions. We thank ABRC (Ohio State University) for providing T-DNA lines seed lines. J.M.E., A.D.N., and J.P.M. are investigators of the National Research Council (CONICET) from Argentina. This work was supported by grants from ANPCyT (PICT2013-003, PICT2014-504, and ICGEB2016 Grant to J.M.E) and from the Mid-career Researcher Program (2015002633) of National Research Foundation and the Next-Generation BioGreen 21 program (The Agricultural Genome Center PJ011195) of the Rural Development Administration to H.-T.C. H.-S.C. was partially supported by the Stadelmann-Lee Scholarship, Seoul National University.

## Author Contributions

S.M., S.P.D.J., H-S.C. and E.M. as co-first authors performed most of the experiments, analysed the data, and wrote the paper. S.M. and S.P.D.-J. isolated and characterized *rboh* mutants, performed ROS detection with the HyPer and H_2_DCF-DA probes, and conducted the pharmacological inhibition and auxin assays. H-S.C. performed the ChIP of ARF5, made ARF5,7,8-overexpressor and pARF5::GUS-GFP lines, and performed Real Time-PCR. E.M. isolated and characterized *per* mutants, measured ROS levels with the HyPer and H_2_DCF-DA probes and carried out pharmacological inhibitions. Y.H. performed the ChIP of RBOHs and PERs. S.M.V. performed ARF5,7,8,19 imaging, analysed the data and commented on the paper contents. C.B., M.L.B., A.A.A, A.D.N., and J.P.M. analysed the data and commented on the paper contents. H-T.C. and J.M.E. designed the research, analysed the data, supervised the project, and wrote the paper. All authors commented on the results and the manuscript. This manuscript has not been published and is not under consideration for publication elsewhere. All the authors have read the manuscript and have approved of this submission.

## REFERENCES

1. B. Menand, et al. An ancient mechanism controls the development of cells with a rooting function in land plants. Science 316, 1477–1480 (2007).

2. S. Datta, H. Prescott, L. Dolan. Intensity of a pulse of RSL4 transcription factor synthesis determines Arabidopsis root hair cell size. Nat. Plants 1, 15138 (2015).

3. E. Slabaugh, M. Held, F. Brandizzi. Control of root hair development in Arabidopsis thaliana by an endoplasmic reticulum anchored member of the R2R3-MYB transcription factor family. Plant J. 67, 395–405 (2011).

4. P. Xu et al. HDG11 upregulates cell-wall-loosening protein genes to promote root elongation in Arabidopsis. J. Exp. Bot. 65, 4285–4295 (2014).

5. K. Yi, B. Menand, E. Bell, L. Dolan. A basic helix-loop-helix transcription factor controls cell growth and size in root hairs. Nat. Genet. 42, 264–267 (2010).

6. J.E. Salazar-Henao, W. Schmidt. An inventory of nutrient-responsive genes in Arabidopsis root hairs. Front. Plant Sci. 7, 237 (2016).

7. Li. Song, H. Yu, X. Che, Y. Jiao, D. Lui. The molecular mechanism of ethylene-mediates root hair development induced by phosphate starvation. Plos Gen. 12(7), e1006194 (2016).

8. T. Guilfoyle, G. Hagen, T. Ulmasov, J. Murfett. How does auxin turn on genes?. Plant Physiol. 118, 341–347 (1998).

9. Fukaki H, Tameda S, Masuda H, Tasaka M Lateral root formation is blocked by a gain-of-function mutation in the SOLITARY-ROOT/IAA14 gene of Arabidopsis. Plant J 29, 153–168 (2002).

10. Bargmann, B.O.R. et al. A map of cell type-specific auxin responses. Molecular Systems Biology, 9, 688. http://doi.org/10.1038/msb.2013.40 (2013).

11. Okushima, Y., et al. Functional genomic analysis of the *AUXIN RESPONSE FACTOR* gene family members in *Arabidopsis thaliana:* Unique and overlapping functions of *ARF7* and *ARF19*. Plant Cell 17, 444–463(2005).

12. J. Foreman et al. Reactive oxygen species produced by NADPH oxidase regulate plant cell growth. Nature 422, 442–446 (2003).

13. J. Wu et al. Spermidine oxidase-derived H2O2 regulates pollen plasma membrane HyPerpolarization-activated Ca2+-permeable channels and pollen tube growth. Plant J. 63, 1042–1053 (2010).

14. Y. Lee, M.C. Rubio, J. Alassimone, N. Geldner. A mechanism for localized lignin deposition in the endodermis. Cell 153, 402–412 (2013).

15. H.-T. Xie, Z.-Y. Wan, S. Li, Y. Zhang. Spatiotemporal Production of Reactive Oxygen Species by NADPH Oxidase Is Critical for Tapetal Programmed Cell Death and Pollen Development in Arabidopsis. Plant Cell 26, 2007–2023 (2014).

16. B. Orman-Ligeza, et al. X. Draye. RBOH-mediated ROS production facilitates lateral root emergence in Arabidopsis. Development 2016 143, 3328–3339 (2016).

17. G. Monshausen, T. Bibikova, M. Messerli, C. Shi, S. Gilroy. Oscillations in extracellular pH and reactive oxygen species modulate tip growth of Arabidopsis root hairs. Proc. Nat. Acad. Sci. USA 104, 20996–21001 (2007).

18. S. Takeda et al. Local positive feedback regulation determines cell shape in root hair cells. Science 319, 1241–1244 (2008).

19. A. Boisson-Dernier et al. ANXUR receptor-like kinases coordinate cell wall integrity with growth at the pollen tube tip via NADPH oxidases. PLoS Biol. 11, e1001719 (2013).

20. F. Passardi, C. Penel, C. Dunand. Performing the paradoxical: how plant peroxidases modify the cell wall. Trends in Plant Sci. 9, 534–540 (2004).

21. D.-W. Kim et al. Functional conservation of a root hair cell-specific cis-element in angiosperms with different root hair distribution patterns. Plant Cell. 18, 2958–2970 (2006)

22. K.S. Brouwer, T. van Valen, D.A. Day, H. Lambers. Hydroxamat estimulated O2 uptake in roots of Pisum sativum and Zea mays, mediated by a peroxidase. Plant Physiol. 82, 236–240 (1986).

23. M. Dynowski, G. Schaaf, D. Loque, O. Moran, U. Ludewig. Plant plasma membrane water channels conduct the signalling molecule H2O2. Biochem. J. 414, 53–61 (2008).

24. S. Tian et al. Plant aquaporin AtPIP1;4 links apoplastic H_2_O_2_ induction to disease immunity pathways. Plant Physiol. 171(3),1635–50 (2016).

25. A. Schlereth et al. MONOPTEROS controls embryonic root initiation by regulating a mobile transcription factor. Nature 464, 913–916. (2010).

26. M. Galli et al. Auxin signaling modules regulate maize inflorescence architecture. Proc. Nat. Acad. Sci. USA 112, 13372–13377 (2015).

27. M.-S. Lee, J.-H. An, H.-T. Cho. Biological and molecular functions of two ear motifs of Arabidopsis IAA7. J. Plant Biol. 59, 24–32(2016).

28. H. Tsukagoshi, W. Busch, P.N. Benfey. Transcriptional regulation of ROS controls transition from proliferation to differentiation in the root. Cell 143, 606–616 (2010).

29. D., Lu, T. Wang, S. Persson, B. Mueller-Roeber, J.H.M. Schippers. Transcriptional control of ROS homeostasis by KUODA1 regulates cell expansion during leaf development. Nat. Commun. 5, 1–9 (2014).

30. P. Vijayakumar, S. Datta, L. Dolan. ROOT HAIR DEFECTIVE SIX-LIKE4 (RSL4) promotes root hair elongation by transcriptionally regulating the expression of genes required for cell growth. New Phytol. doi:10.1111/nph.14095 (2016).

31. Y. Hwang, H.S. Choi, H.M. Cho, H.T. Cho. Tracheophytes Contain Conserved Orthologs of a Basic Helix-loop-helix Transcription Factor to Modulate ROOT HAIR SPECIFIC Genes. Plant Cel. doi:10.1105/tpc.16.00732 (2017).

32. K. Sundaravelpandian, N.N. Chandrika, W. Schmidt. PFT1-controlled ROS balance is critical for multiple stages of root hair development in Arabidopsis. New Phytol. 197(1), 151–61 (2013).

33. B.W. Yun et al. S-nitrosylation of NADPH oxidase regulates cell death in plant immunity. Nature 478, 264–268 (2011).

34. J. Nestler et al. Roothairless5, which functions in maize (Zea mays L.) root hair initiation and elongation encodes a monocot-specific NADPH oxidase. Plant J. 79, 729–740 (2014).

35. T. Kwon et al. Transcriptional response of Arabidopsis seedlings during spaceflight reveals peroxidase and cell wall remodeling genes associated with root hair development. Am J Bot. 102, 21–35. (2015).

36. C. Dunand, M. Crèvecoeur, C. Penel. Distribution of superoxide and hydrogen peroxide in Arabidopsis root and their influence on root development: possible interaction with peroxidases. New Phytol. 174, 332–341(2007).

37. E.W. Miller, B.C. Dickinson, C.J. Chang. Aquaporin-3 mediates hydrogen peroxide uptake to regulate downstream intracellular signaling. Proc. Nat. Acad. Sci. USA 107, 15681–15686. (2010).

